# Computational Psychiatry Research Map (CPSYMAP): a New Database for Visualizing Research Papers

**DOI:** 10.1101/2020.06.30.181198

**Authors:** Ayaka Kato, Yoshihiko Kunisato, Kentaro Katahira, Tsukasa Okimura, Yuichi Yamashita

## Abstract

The field of computational psychiatry is growing in prominence along with the recent advances in computational neuroscience, machine learning, and the cumulative scientific evidence of psychiatric disorders. Computational approaches provide an understanding of disorders using psychological and neuroscience terms and help to determine treatment protocols based on high-dimensional data. However, the multidisciplinary nature of this field seems to limit the development of computational psychiatry studies. Computational psychiatry combines knowledge from neuroscience, psychiatry, and computation; thus, there is an emerging need for a platform to integrate and coordinate these perspectives. In this study, we developed a new database for visualizing research papers as a two-dimensional “map” called the Computational Psychiatry Research Map (CPSYMAP). This map shows the distribution of papers along neuroscientific, psychiatric, and computational dimensions to enable anyone to find niche research and deepen their understanding of the field.

## 1. Introduction

The understanding of psychiatric disorders is based on multiple interacting levels, from genetics to cells, neural circuits, cognition, behavior, and the surrounding environment. Recent advances in computational methodology and cumulative biological and psychological evidence have helped to handle high-dimensional data to understand and treat psychiatric disorders [1-5]. Bringing mathematical methodologies and theoretical frameworks to psychiatric research is expected to have a central role in the development of treatments and preventive strategies; the use of such methods constitutes an area of research called computational psychiatry (CPSY), which has come to be treated as an important psychiatry discipline. The development of this field has been reviewed in many papers from the clinical and computational neuroscientific perspective [6-11]. This is a highly multidisciplinary field that requires researchers from a variety of backgrounds ranging across neuroscience, psychiatry, and mathematical modeling. Therefore, to accelerate CPSY research, it is crucial to link each CPSY study to the knowledge derived different fields, including accumulated evidence of neuroscience, traditional psychiatry, and computational techniques.

### 1.1 Link to Neuroscience

Traditionally, mental disorders have been diagnosed based on patients’ self-reporting of subjective experiences and behavioral observation by clinicians; disorders are conceptualized by accumulating those observations. While this view provides benefits such as reliability of diagnosis, this approach presents challenges such as lack of biological plausibility and predictive validity [12]. The National Institute of Mental Health (NIMH) has developed the Research Domain Criteria (RDoC) framework to systematize psychiatric research based on the findings of behavioral neuroscience that is not bound by conventional disease categories [13-14]. Moreover, they have provided the RDoC matrix, which has integrated many levels of information. The observable basic component functions based on behavioral neuroscience are defined as constructs. Psychiatric disorders are considered on a spectrum from normal to abnormal states of constructs and related constructs are organized into a domain. In addition, they enumerated the variables of the unit of analysis from micro to macroscopic levels. However, the correspondence between the contents of each CPSY research paper and RDoC is not mentioned in many cases. Linking RDoC to each study based on the accumulated neuroscience findings is essential to enhance the significance of CPSY research, particularly with regard to showing the expected role of CPSY research in connecting the knowledge of different aspects using mathematical models.

### 1.2 Link to Traditional Psychiatry

Since RDoC does not cover the traditional taxonomy and symptomatology of mental disorders, linking CPSY studies to RDoC is not sufficient. As mentioned, traditional psychiatric symptomatology has contributed significantly to the research and diagnostic classification criteria because of the inter-rater reliability [15], which has been accumulated in the fifth edition of the Diagnostic and Statistical Manual of Mental Disorders (DSM-5) [16,17]. Thus, an integration of the terms of psychiatric symptomatology and neuroscientific knowledge defined in the RDoC is needed. However, it is difficult for researchers outside the domain of psychiatry to obtain an overview of which symptoms are important and what should be studied.

### 1.3 Link to Computation

In CPSY research, various powerful computational tools and conceptual frameworks are used to analyze and manipulate high-dimensional, multimodal data sets, including clinical, genetic, behavioral, neuroimaging, and other data types obtained by animal models, human experiments, clinical reports, and simulations. In this domain, there are different cultures, such as the theory-driven vs. data-driven approaches [11]. The theory-driven approach models the information processing of perception and cognition in the brain based on the computational framework. The data-driven approach applies machine learning methods to large-scale data on mental disorder, then clusters the data and develops discriminatory models. Additionally, various types of models (e.g., reinforcement learning, neural networks, and biophysical models) have been utilized in this field. It is, therefore, necessary to organize the methodologies, data, theories, and models that are used to accelerate the research.

### 1.4 Database to integrate the three perspectives

Due to the cumulative efforts in the fields of neuroscience, psychiatry, and mathematical modeling, CPSY research could have a central role in the rational development of psychological pathology, treatments, ontologies, and preventive strategies. However, insufficient connections between the perspectives of neuroscience, psychiatry, and computational models may hinder the extension of research. To enable further development in the field, it is necessary to organize individual studies from the following perspectives: 1) Connection to neuroscience: What cognitive functions and units of analysis, such as behavior and circuitry, are targeted?, (2) Connection to psychiatry: Which mental disorders and symptoms are being addressed and, which are not being addressed?, (3) Connection to computation: What methodologies, data, theories, and models are used?

To address this issue, we developed a database and created an environment wherein anyone can search, register, and obtain an overview of CPSY research as a web application. In this report, a CPSY Research Map (CPSYMAP) (accessible online at https://ncnp-cpsy-rmap.web.app/). [18] is described, which is an online database that tags CPSY papers using the terminology of neuroscience, psychiatry, and computation to help researchers organize and visualize the status of research areas along with the tags on a two-dimensional map.

## 2. Method

### 2.1 Database architecture

CPSYMAP is accessed through a public website at https://ncnp-cpsy-rmap.web.app. The database was built using Google firebase [19] and the source code was written in Javascript. The core of CPSYMAP is a database consisting of two components: information about the individual papers and the tags based on neuroscientific, psychiatric, and computational knowledge. The information about the papers is taken from API Crossref [20]. The data points stored are the title of the paper, the names of authors, the citation count, digital object identifier (DOI), data type, and abstract.

### 2.2 Tags used in the database

Individual CPSY papers are tagged according to the categories of neuroscience, psychiatry, and computation in order to characterize and visualize their relationships from the three perspectives described above; lists of all tags are shown in Supplementary Table. 1.

From the neuroscientific perspective, the RDoC domain, construct, and subconstruct are provided as options showing which cognitive functions are targeted. To arrange the papers by the unit of analysis, users can choose the tags from genes, molecules, cells, neural circuits, physiology, behaviors, self-reports, and paradigms.

To visualize the mental disorders and symptoms are being addressed and those not being addressed, we provide options from both the DSM-5 and symptomatology. The DSM-5 is the authoritative list of mental disorders proposed by the American Psychiatric Association [16]. Psychiatric symptomatology is a method of psychiatry attempting to describe and reconstruct the characteristics of patients’ experiences from the aspects of psychology and phenomenology. We are referencing the lexicon of psychiatric and mental health terms (WHO) [17].

To arrange the different types of computational frameworks, we provide options called experimental design. In this category, we defined seven tags including Generative model, Model-fitting, Simulated lesion, Classification/Discrimination, Clustering, Prediction of disease states and Others. The definitions are provided in the database [1] and in Table 1.

**Table 1.**
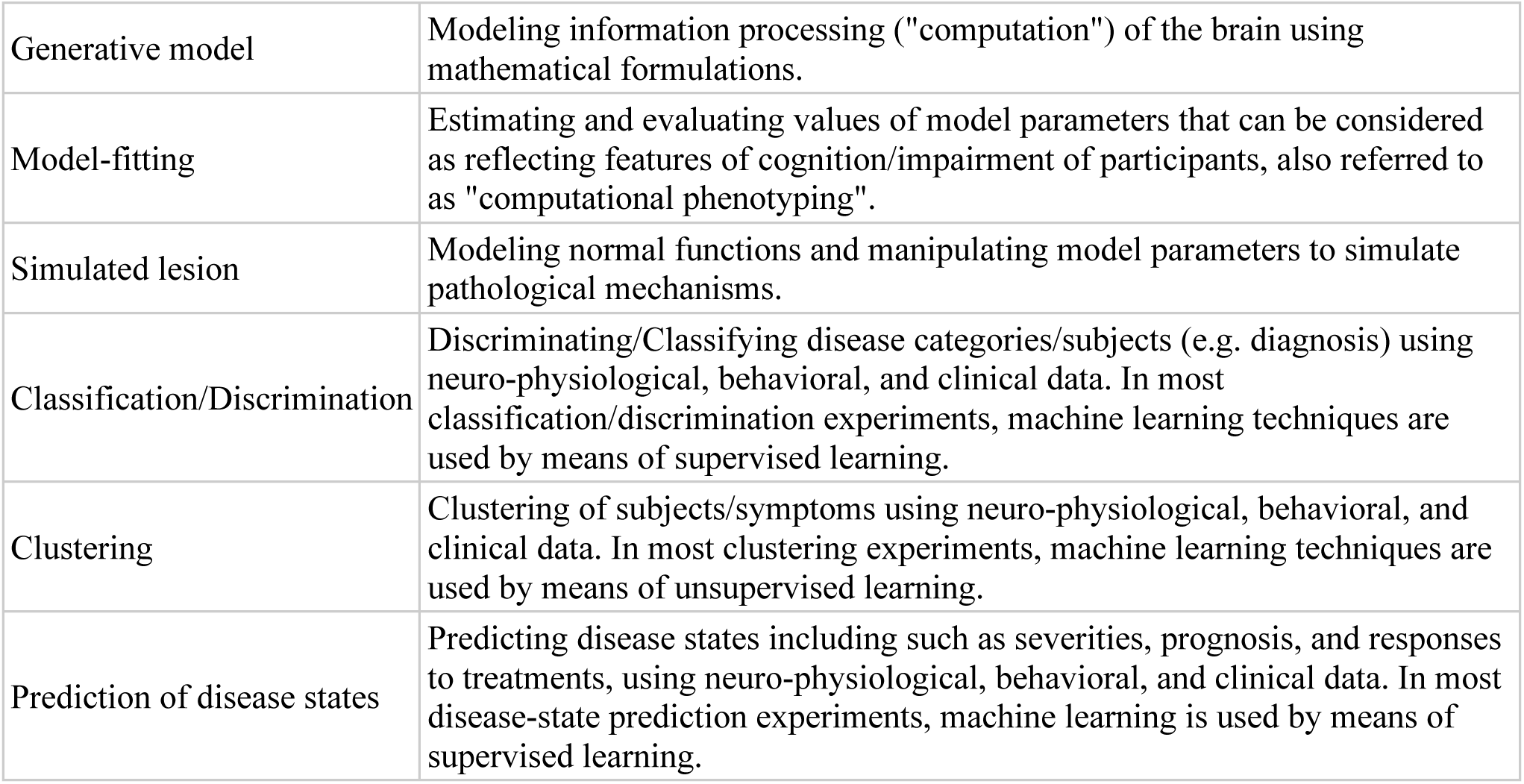
Definition of the experimental design.

The data type can be selected from animal data, human data, simulation, and literature review. Although we could not cover all the models used in this field, the tags the for models are also situated. For the model names, 13 tags are prepared and 5 models (Reinforcement learning, Biophysical model, Neural network model, Bayes and Regression model) are shown on the axis.

### 2.3 Organization

CPSYMAP provides a simple graphical user interface to create a web application environment where anyone can search, register and, overview CPSY research. In addition, users can easily select and tag the scientific papers they want to add.

### 2.4 Reliability of CPSYMAP

This is a community augmented approach, meaning that CPSYMAP is publicly editable. Thus, like other crowd-sourced projects, the registered information is not guaranteed to be accurate. We aimed to avoid flooding the CPSYMAP with inaccurate information and making it less reliable by implementing the following measures. First, if a user adds a tagged paper, they can review and edit the related tag information at any time. The members in Computational Psychiatry Colloquium, a volunteer group for the study of CPSY, occasionally add papers and check the information registered in the database. We have also implemented a simple reporting system, called “Edit request,” on the individual page of every paper so that other users may flag mistakes or make suggestions regarding the tags added on papers by other users. Editors of the Computational Psychiatry Colloquium group validate these requests and make appropriate changes to the database.

## 3. Results

### 3.1 Visualizing, Filtering, and Sorting

This database organizes and visualizes the status of the research areas along the tags on a two-dimensional map. In the Heatmap section, the green color shading indicates areas with many papers and areas with few papers (Figure 1).

**Figure 1.**
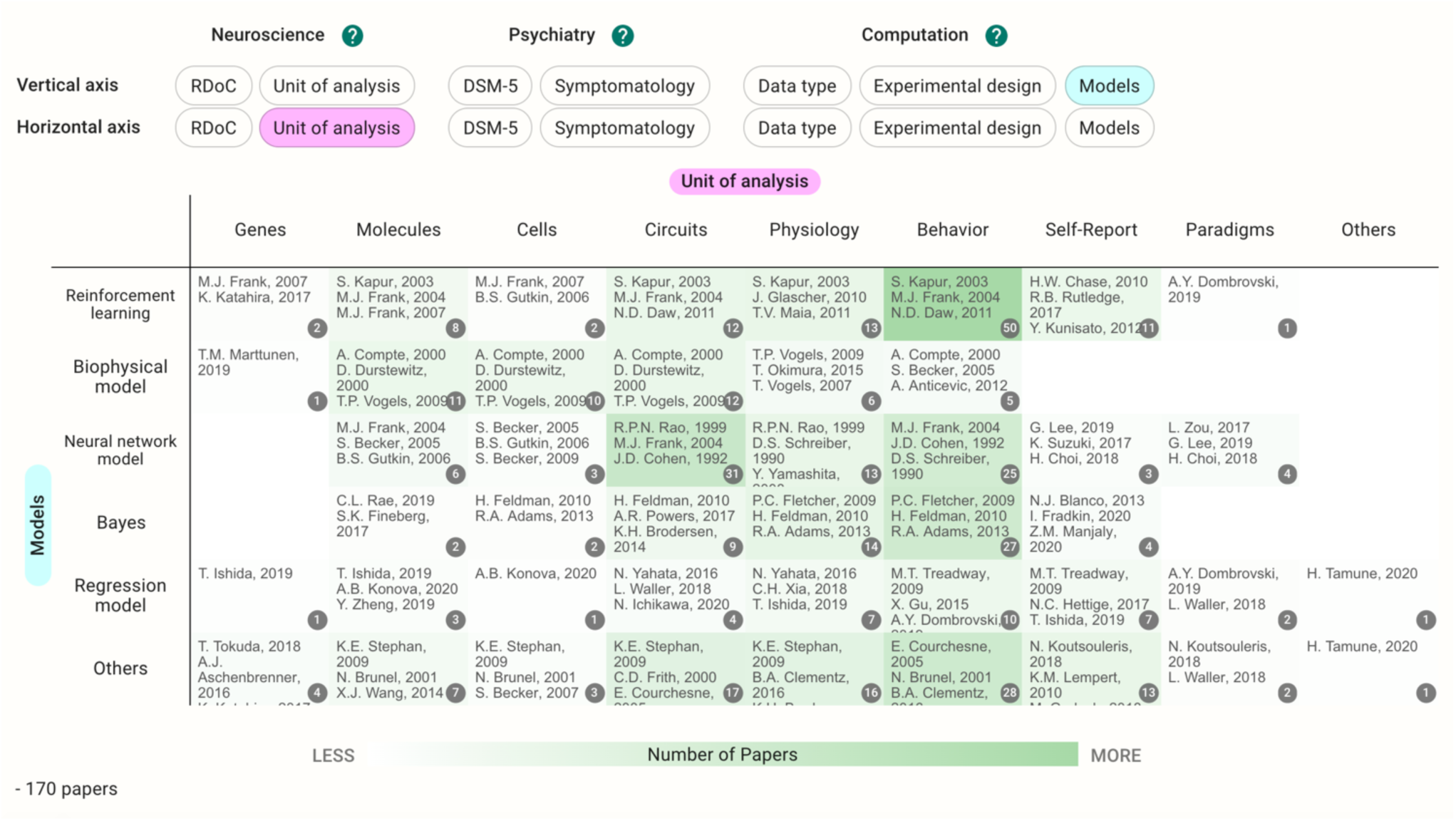
Example view of the heatmap section.

Each cell contains the papers with tags specified in the vertical and horizontal axes. The users can choose the vertical and horizontal axes they wish to view. Clicking the cell takes users to the Pickup Table. This section allows users to view the paper list in the cell and sort the tags in ascending or descending order. The details of the paper appear when users choose a paper from the table. The filter function can be used to further narrow the contents in the map.

### 3.2 One-dimensional analysis of registered papers

Using these functions, we introduced a preliminary notice from the database. When accessed on 10th June 2020, there were 170 papers in total. If users choose the same category on both the horizontal and vertical axes, they can see the distribution of the papers on a single axis (Figure 2). Since we allow multiple selections of tags during the registration, there are some papers outside of the diagonal cells. In the case of the data type, for example, 57 papers used the simulation as their main result and 110 papers reported experiments with human participants; 13 papers dealt with both simulations and human experiments.

**Figure 2.**
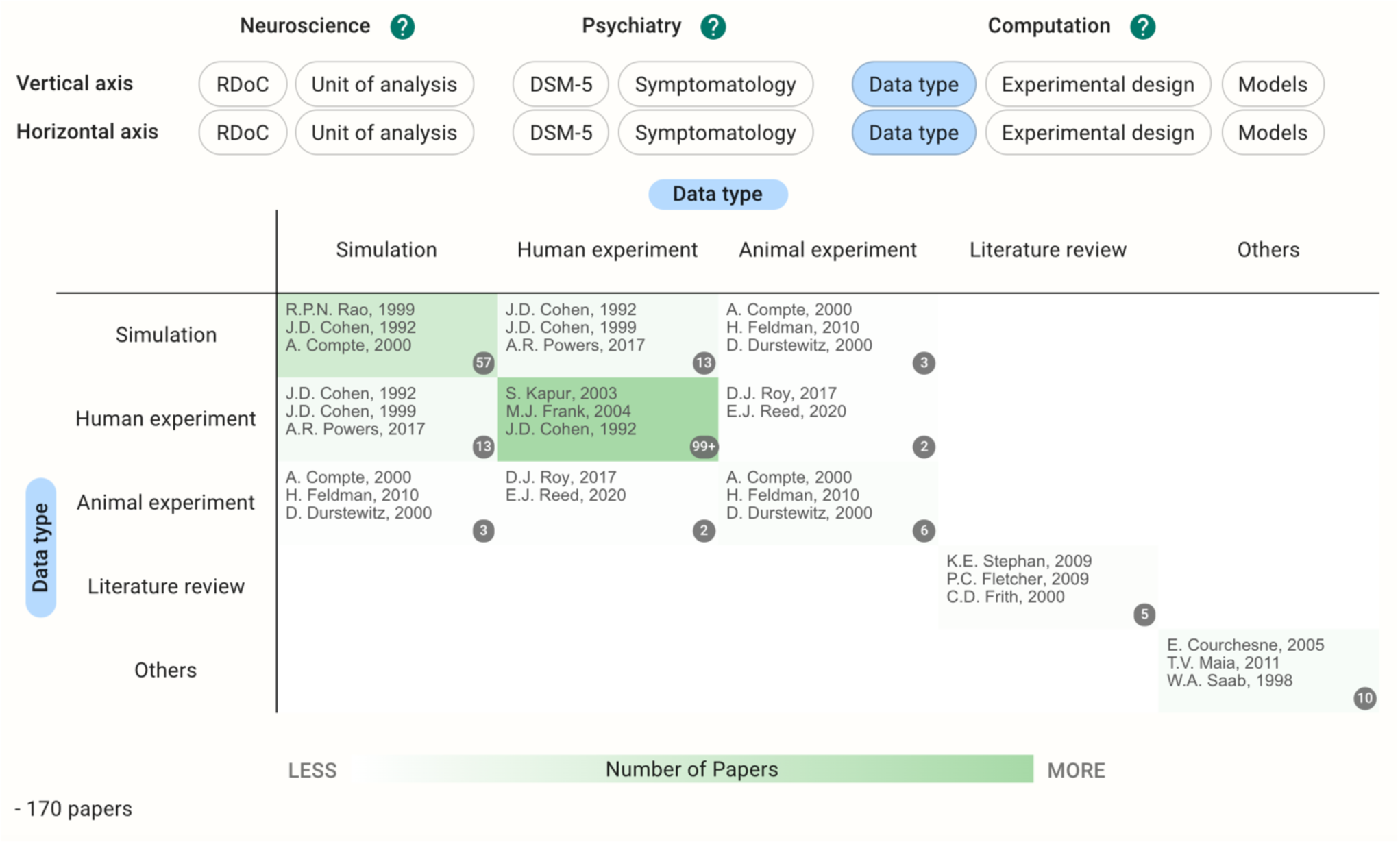
Heatmap with data type in both horizontal and vertical axes.

Although this trend could be temporary, the most popular diseases discussed in this database were schizophrenia spectrum and other psychotic disorders (68 papers, Figure 3). The column called “Others” included papers targeting 13 other disorders and normal cognitive functions that were tagged as cognitive processes at the registration process (Figure 3, bottom). Regarding the experimental design from the computational perspective, both generative models (62 papers) and model fitting (60 papers) were frequently used in the registered CPSY papers (Figure 4). However, all designs were used to some extent, showing the variety of the methodology in the field.

**Figure 3.**
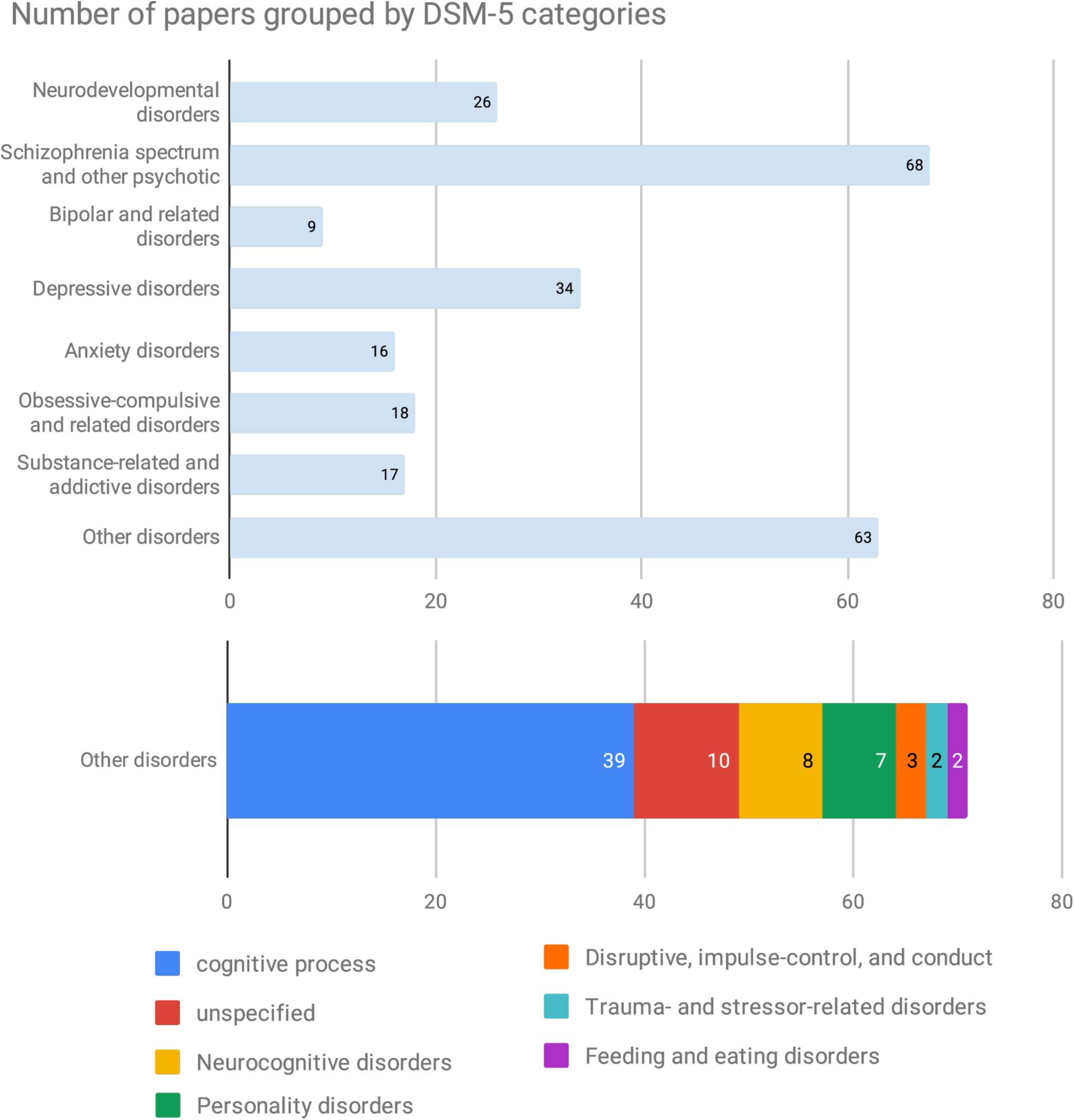
Number of papers grouped by DSM-5 categories.

**Figure 4.**
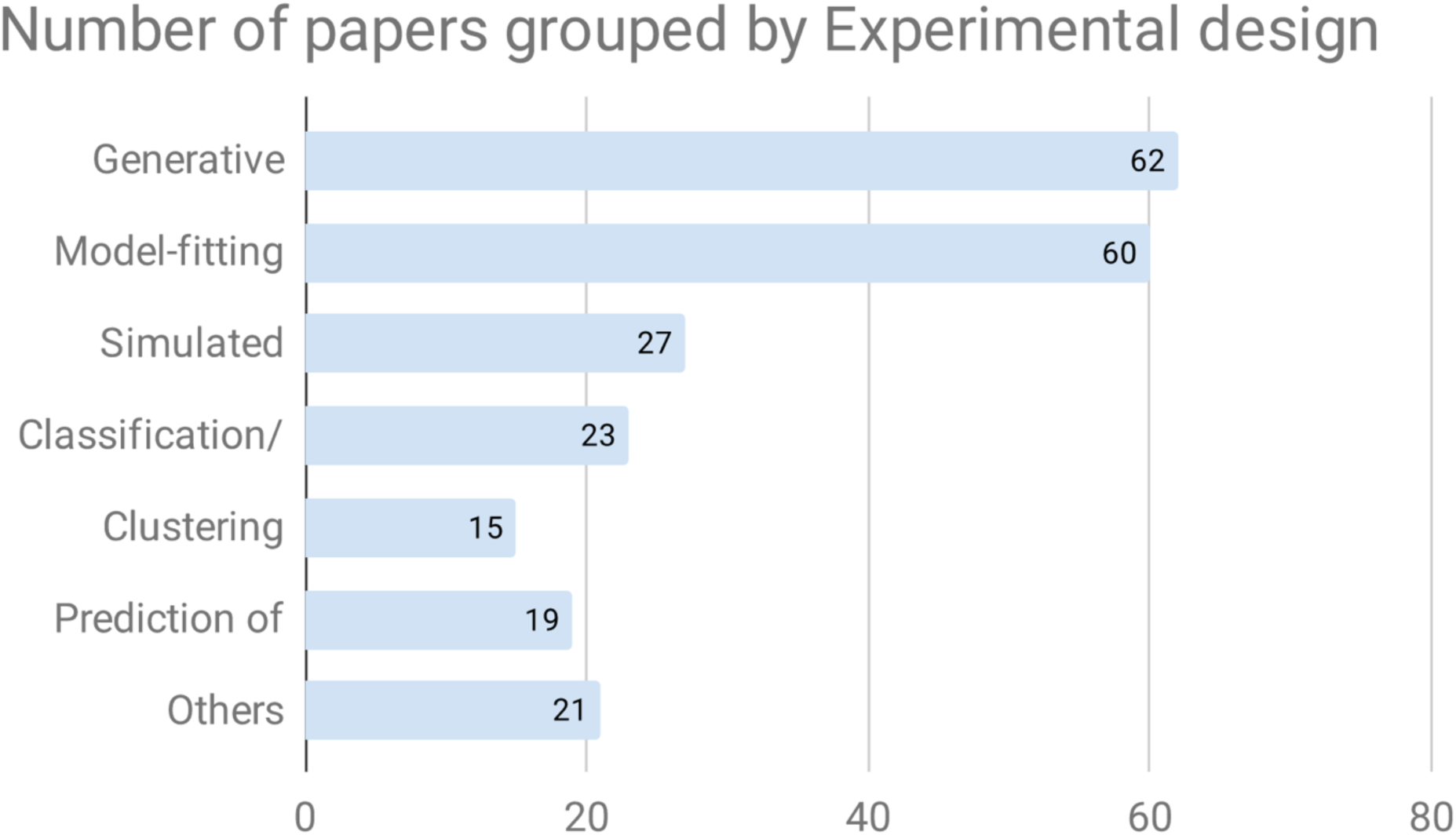
Number of papers grouped by experimental design.

Regarding the RDoC domain, the cognitive system was the term most frequently tagged to papers (82 papers). By using the pickup table and filter function, users can shortlist the papers. Within the constructs of the cognitive system, cognitive control was tagged to 34 papers, while working memory and perception were studied in 21 papers.

### 3.3 Two to three-dimensional analysis of the registered papers

If the unit of analysis and models are chosen for the axes in the current database, the cells with the largest number of papers are on behavior crossed with reinforcement learning (50 papers, Figure 1). The papers included in the cell dealt with a wide range of disorders such as schizophrenia (10 papers) and substance use disorders (10 papers); however, depressive disorders were the most prevalent (16 papers).

In cases where the target disease is already determined, the users can see the different landscapes by choosing the disease in the filter and perusing the Heatmap section. Before filtering by the DSM-5 category, the reinforcement learning crossed with model fitting (35 papers) is the most common methodology (Figure 5); however, the cell with the neural network model crossed with the generative model has the largest number of the papers (12 papers) after the filtering by schizophrenia (Figure 6).

**Figure 5.**
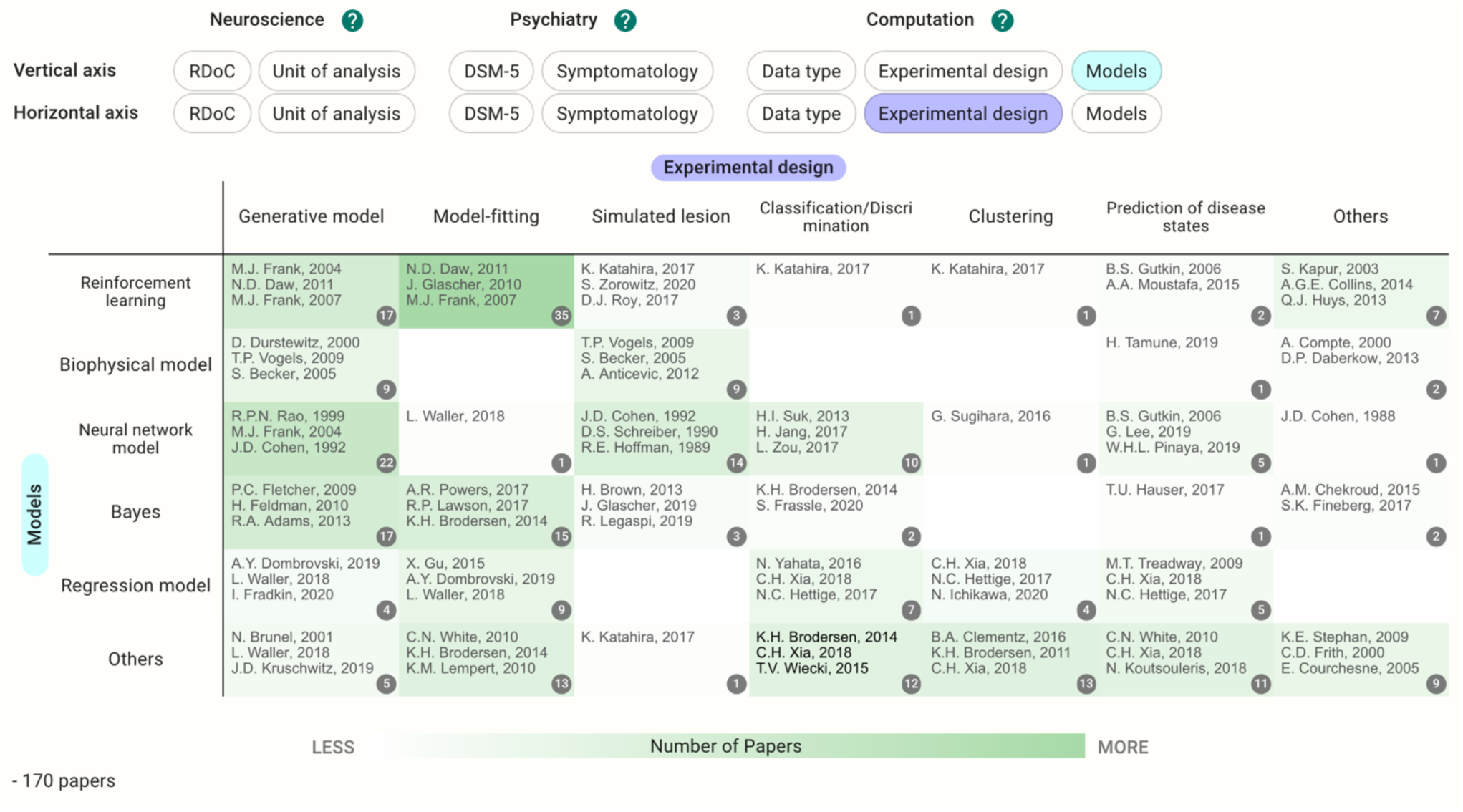
Heatmap with models and experimental design as the vertical and horizontal axes, respectively.

**Figure 6.**
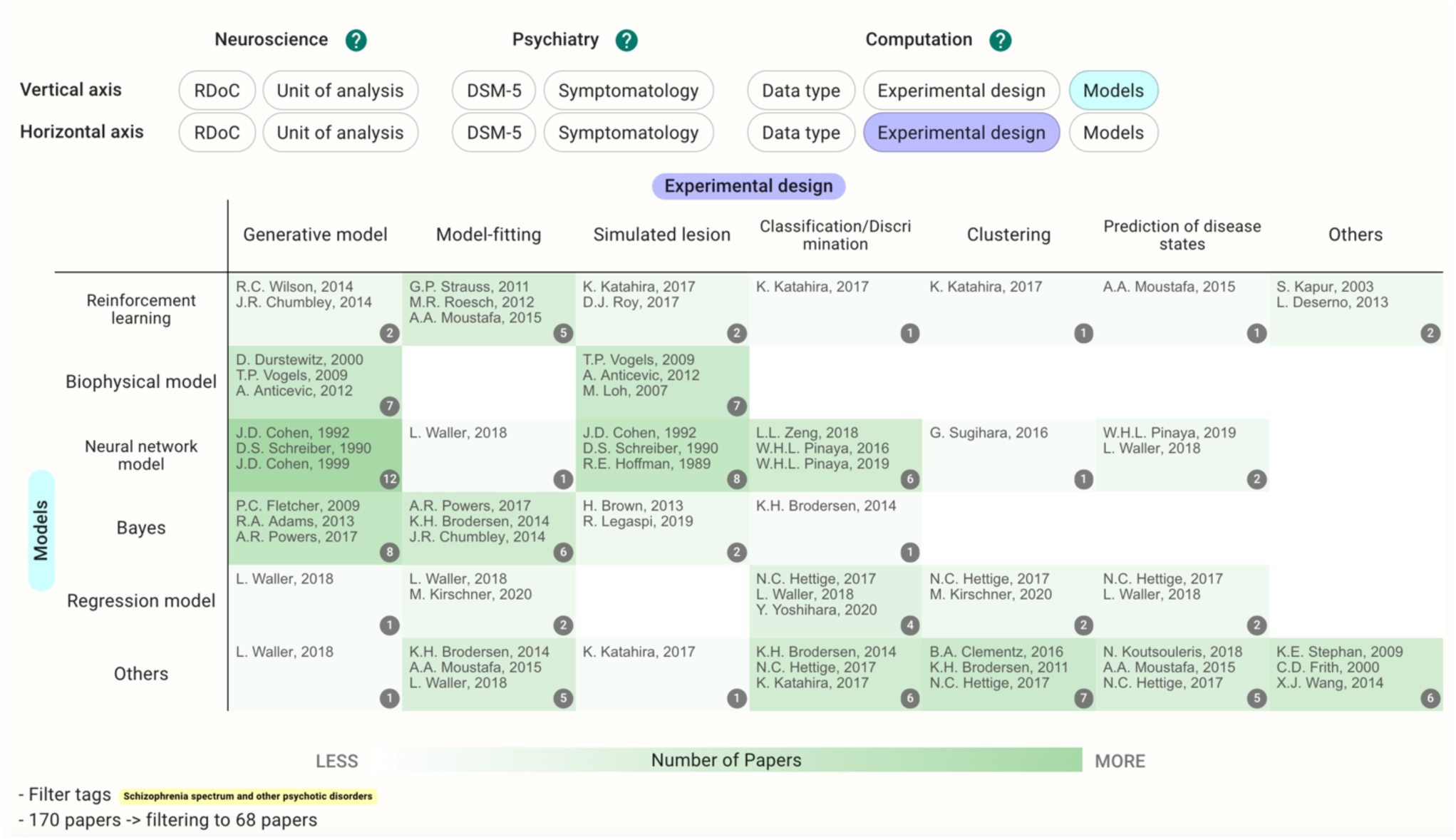
Heatmap with models and experimental design as the vertical and horizontal axes, respectively, filtered by DSM-5 category, schizophrenia spectrum, and other psychotic disorders.

## 4. Discussion

We developed CPSYMAP for visualizing research papers on a two-dimensional map and to provide an interactive environment where anyone can search, register, and obtain an overview of CPSY research. Further, we described the details and importance of the perspectives of neuroscientific, psychiatric, and computational methods and how we implemented those dimensions in this database. Finally, we showed some preliminary analysis of the registered papers in the database.

This is the first database of research papers in the field of CPSY. In addition, the CPSYMAP provides the benefit of presenting new ways of organizing papers, such as expressing distributions with the use of colors in maps and switching axes. It is designed in a way such that it can be updated and augmented with the mutual efforts of the community. This can be a new form of reviewing research articles in the sense that the database can help researchers design new research plans and organize previously published works.

However, other approaches to organize and visualize research papers are also available. For instance, Microsoft Academia is a semantic search engine for academic research and is not keyword-based [21]. It provides a sophisticated user interface to visualize the keywords of papers and organize the academic knowledge including relationships between authors in a variety of research fields. However, since the scope of this engine includes the entire academia, it may not cover the granularity we aimed to achieve in our database. A notable advantage of our database is that it attempts to cover the entire field of CPSY while ensuring a fine granularity and organized dimensionality (neuroscience, psychiatry, and computation).

In addition, the significance of systematic evidence maps in several fields, including epidemiology [22], environmental science [23], and psychiatry [24, 25, 26], is increasing. These maps provide a comprehensive overview of the evidence with visual depictions such as figures or tables in most cases [27]. CPSYMAP can be considered as one form of an evidence map. However, a unique feature of our database it can be updated to include all the newly published papers in this field. This allows for an open and community-augmented approach, which is expected to promote the field of computational psychiatry.

On the contrary, there are some limitations to this database. Currently, the number and categories of registered papers are limited. However, this can be resolved as the community begins to assist us with registration of additional research papers and as more computational psychiatrists become familiar with our database. We plan to use natural language processing to recommend potential papers that are suitable for registration in our database and suggest tags to the individual papers.

The second limitation is the potential for credibility issues with tagging. Specifically, there may be disagreements and differing views on tagging among the users and contributors in the community. At present, users are asked to identify problems using the edit request function. However, we are also considering disclosing the discussion process regarding which tags are appropriate to be placed on a bulletin board.

## 5. Conclusion

We expect that the CPSYMAP database will provide a better perspective for non-specialists regarding topics that are not explored often and that are important in the CPSY field. This may attract more experts from other fields such as informatics, physics, and engineering, as well as more experts from psychiatry and neuroscience to CPSY research.

Moreover, the user interface allows researchers to understand CPSY papers with a multidimensional view, which we hope will assist with the development of computational methods and theoretical frameworks to establish links between traditional psychiatry and neuroscience. This can contribute to the integration of different methods, especially the interactive and cyclical development of both the theory-driven and data-driven approaches. By building a platform for organizing and storing information, we hope that the database contributes to the efficiency of research and vitalization of the field.

## 7. Author Contributions

AK and YY conceived the study and wrote the manuscript. TO, YK, and KK advised on the system design of the database. They also reviewed and commented on the manuscript. All authors contributed to the preparation of the article and approved the final version.

## 9. Acknowledgments

We wish to thank Mr. Yuji Kawase for developing the database. We also wish to thank Mr. Chris Salzberg and Editage for English editing, and thank Dr. Hiroshi Yamakawa for stimulating discussions. Finally, this database is also supported by the members of the computational psychiatry colloquium in Japan. We appreciate their suggestions for the user interface and efforts to register papers in the database.

